# Improved G-AgarTrap: A highly efficient transformation method for intact gemmalings of the liverwort *Marchantia polymorpha*

**DOI:** 10.1101/329839

**Authors:** Shoko Tsuboyama, Satoko Nonaka, Hiroshi Ezura, Yutaka Kodama

## Abstract

Liverworts are key species for studies of plant evolution, occupying a basal position among the land plants. *Marchantia polymorpha* has emerged as a highly studied model liverwort, and many relevant techniques, including genetic transformation, have been established for this species. *Agrobacterium*-mediated transformation is widely used in many plant species because of its low cost. Recently, we developed a simplified *Agrobacterium*-mediated method for transforming *M. polymorpha*, known as AgarTrap (a<u>gar</u>-utilized transformation with pouring solutions). The AgarTrap procedure, which involves culturing the liverwort tissue in various solutions on a single solid medium, yields up to a hundred independent transformants. AgarTrap is a simple procedure, requiring minimal expertise, cost, and time.

Here, we investigated four factors that influence AgarTrap transformation efficiency: (1) humidity, (2) surfactant in the transformation buffer, (3) *Agrobacterium* strain, and (4) light/dark condition. We adapted the AgarTrap protocol for transforming intact gemmalings, achieving an exceptionally high transformation efficiency of 97%. The improved AgarTrap method will enhance the molecular biological study of *M. polymorpha*. The present study also provides new possibilities for improving transformation techniques for a variety of plant species.

## Introduction

*Marchantia polymorpha* is a dioecious liverwort, the sister group to all other land plants^1^. This species has therefore been extensively studied to enhance our understanding of land plant evolution, with research focusing on its taxonomy, development, and physiology; furthermore, its nuclear, chloroplast, and mitochondrial genomes have all been sequenced^2–6^. The rapidly expanding *M. polymorpha* research community has recently developed various molecular biology techniques to study this key species, including particle bombardment-and *Agrobacterium*-mediated transformation, plastid transformation, homologous recombination-mediated gene targeting, and TALEN-and CRISPR/Cas9-mediated genome editing^7–16^.

*Agrobacterium*-mediated transformation is widely used for many plant species because it does not require any expensive equipment^17^. This technique involves three steps: (1) preparation of plant material, (2) co-culture of the material with *Agrobacterium tumefaciens* containing a recombinant transfer DNA (T-DNA), and (3) antibiotic selection of transgenic cells. During the co-culture step, T-DNA is transferred from the *Agrobacterium* into the plant cell, where it is integrated into the genome to facilitate the expression of its constituent genes. Previous studies have determined that the co-culture conditions are the most important aspect of transformation efficiency, with the *Agrobacterium* strain used, duration of co-culture, *Agrobacterium* density, temperature, co-culture medium, and surfactants used having the greatest impact^18–22^.

Recently, we developed a simplified *Agrobacterium*-mediated transformation method for *M. polymorpha*, which we named AgarTrap (agar-utilized transformation with pouring solutions)^23–25^. Like the general *Agrobacterium*-mediated transformation procedure, AgarTrap consists of three steps: (1) pre-culture of *M. polymorpha* tissue, (2) co-culture of the tissue with *Agrobacterium* containing recombinant T-DNA, and (3) selection of transgenic cells. A unique feature of AgarTrap is that none of these steps require a liquid medium culture; rather, the appropriate solutions are simply poured onto the solid medium in a single Petri dish (Fig. 1)^23–25^. We previously developed and optimized AgarTrap for use with *M. polymorpha* sporelings (S-AgarTrap), intact gemmae/gemmalings (G-AgarTrap), and pieces of mature thallus (T-AgarTrap), achieving transformation efficiencies of approximately 20%, 60%, and 70%, respectively^23–25^. Despite its low transformation efficiency, S-AgarTrap results in numerous transformants, because spores are produced abundantly, rendering it suitable for the large-scale production of transformants (e.g., T-DNA insertion mutants)^23^. However, because spores are produced by sexual reproduction, S-AgarTrap transformants do not have a uniform genetic background. G-AgarTrap can be used to produce transformants in a genetically uniform background, because the gemmae develop from single cells asexually generated within the gemma cup on a mature thallus^24,26^. Similarly, T-AgarTrap results in transformants with uniform genetic backgrounds, because the cells are obtained from mature thalli^25^; however, fewer individual transformants were obtained using T-AgarTrap than G-AgarTrap despite their respective transformation efficiencies, because the pieces of mature thallus were larger than the gemmae and fewer could be included in a single Petri dish. Thus, of these three methods, G-AgarTrap appears to be the best choice for producing transgenic *M. polymorpha*; however, because the transformation efficiency of G-AgarTrap was relatively low (approximately 60%), this approach needed improvement. As the co-culture step is the most critical for efficient transformation^18,22^, the transformation efficiency of G-AgarTrap would likely be improved by optimizing this step.

**Fig. 1.**
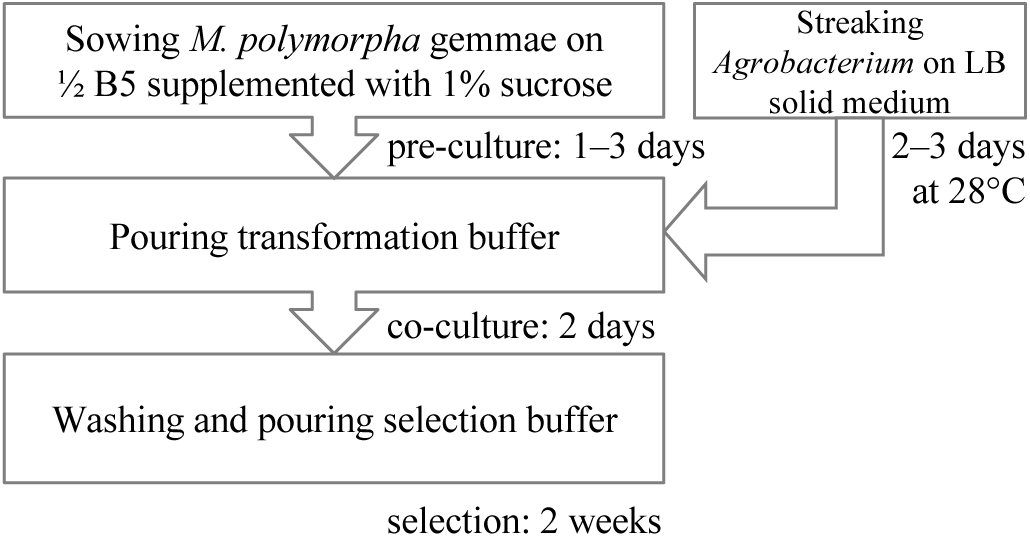
Flowchart of G-AgarTrap. Step I: Pre-culture of *M. polymorpha* gemmae/gemmalings on ½ B5 supplemented with 1% sucrose, and *Agrobacterium* on LB agar medium. Step II: Co-culture of *M. polymorpha* gemmalings with *Agrobacterium* on ½ B5 supplemented with 1% sucrose. Step III: Washing of *M. polymorpha* gemmalings and selection of transgenic cells on ½ B5 supplemented with 1% sucrose.

In our previous study, we optimized several factors of AgarTrap transformation, including the pre-culture period of *M. polymorpha* tissue, the duration of co-culture, *Agrobacterium* density (OD_600_ in transformation buffer), acetosyringone concentration in the transformation buffer, medium composition, and *Agrobacterium* culture conditions^23–25^. In the present study, we investigated four additional co-culture factors: (1) humidity, (2) surfactant in the transformation buffer, (3) *Agrobacterium* strain, and (4) light/dark condition. We also fine-tuned the pre-culture period, ultimately achieving an exceptionally high transformation efficiency for the G-AgarTrap procedure, of nearly 100%.

## Results

### Humidity conditions during co-culture

In our previous study, gemmalings (BC3-38) were pre-cultured for one day and co-cultured with *Agrobacterium* for three days on ½ B5 medium supplemented with 1% sucrose, which resulted in a median transformation efficiency of 57.0% (mean: 59.2%) (Fig. 2a)^24^. Permeable microporous tape was used to seal the Petri dishes containing the solid medium; therefore, the humidity to which the plants were exposed depended on the humidity of the culture room. The humidity of the culture room was kept at approximately 40% with a humidifier, as in our previous study^24^. In the present study, we tested whether humidity differences in the co-culture step influence transformation efficiency. Without the humidifier, the humidity in the culture room decreased to approximately 20%. When gemmalings were co-cultured with *Agrobacterium* at 20% humidity, the median transformation efficiency was decreased by 8.1%, and the mean efficiency decreased by 10.5% (Fig. 2a, see also Supplementary Table S1). These results suggested that higher humidities during the co-culture step increase transformation efficiency; however, it can be challenging to control the humidity in culture rooms precisely, because humidity fluctuates depending on the location and/or season.

**Fig. 2.**
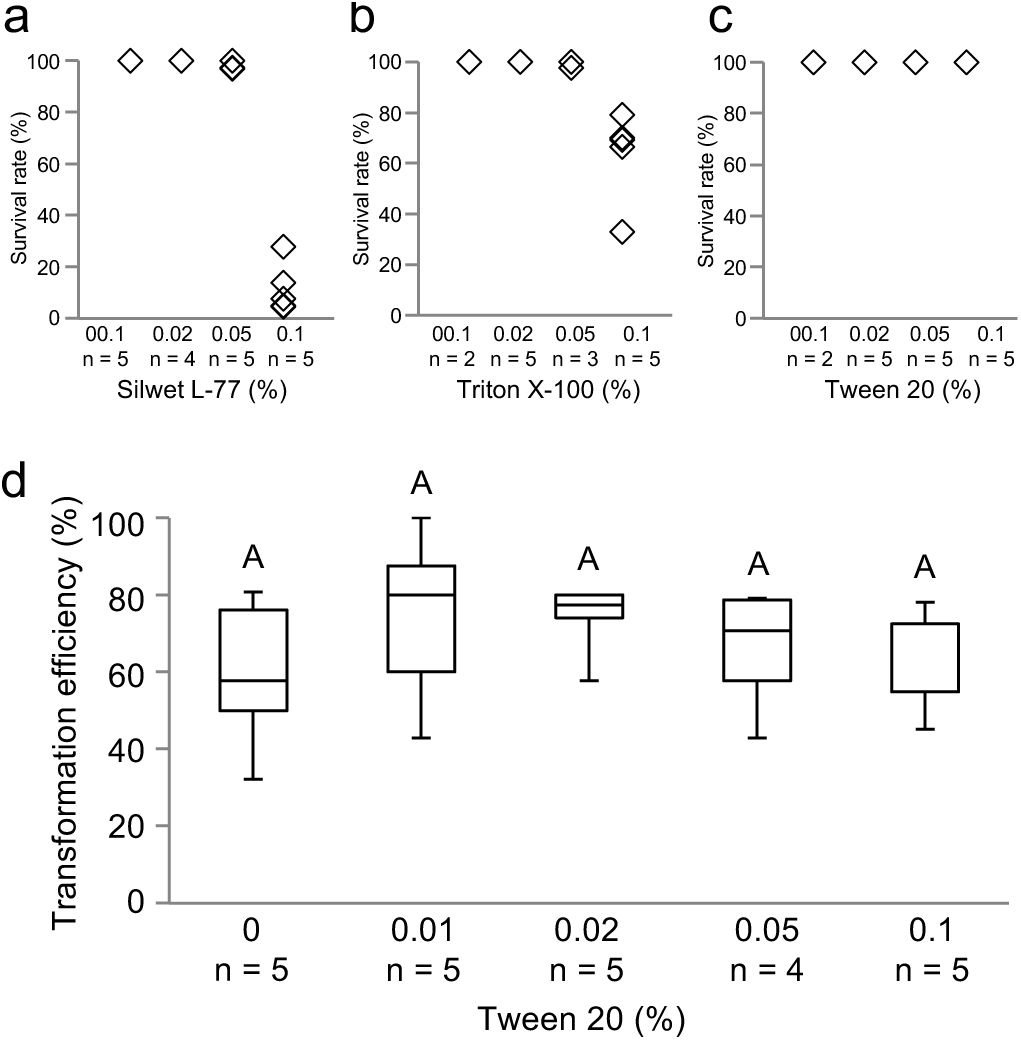
Effect of sealing culture dishes with Parafilm on transformation efficiency. (a) Comparison among the use of microporous tape to seal Petri dishes during co-culture in a culture room at approximately 40% and 20% humidity, and the use of Parafilm to seal Petri dish during co-culture in a culture room at approximately 20% humidity. For both examinations using microporous tape, gemmalings were subjected to a one-day pre-culture before a three-day co-culture. For examinations using Parafilm, gemmalings were subjected a one-day pre-culture before a two-day co-culture. All examinations were performed with *Agrobacterium* strain GV2260 under light. Different letters indicate a significant difference (Tukey’s test; P<0.05). *These raw data were reported in Tsuboyama-Tanaka & Kodama 2015. (b) Effect of the duration of gemmaling pre-culture prior to the use of Petri dishes sealed with Parafilm during the co-culture. All examinations were performed after a two-day co-culture with *Agrobacterium* strain GV2260 under light. Different letters indicate a significant difference (Tukey-Kramer’s test; P<0.05).

To maintain a high humidity in the Petri dishes during co-culture, we sealed the dishes with Parafilm, which is more airtight than microporous tape. When using Parafilm, almost all gemmalings co-cultured for three days suffered from an overgrowth of *Agrobacterium* such as Supplementary Fig. S1, suggesting that the growth of this bacterium is enhanced by high humidity. Since it was difficult to completely eliminate the bacteria in the subsequent selection step when they were overgrown, the co-culture period was shortened to two days when using Parafilm, which increased the median transformation efficiency to 62.3% (mean: 59.6%) in the 20% humidity condition (Fig. 2a, see also Supplementary Table S1). These results indicate that the high humidity in Parafilm-sealed Petri dishes during the co-culture step increases the transformation efficiency, while shortening the required duration of the co-culture period from three days (at 40% humidity when sealed with microporous tape) to two days.

Next, we investigated the pre-culture period of gemmae/gemmalings required when sealing the dishes with Parafilm during the co-culture step. The gemmalings were pre-cultured for 0, 1, 2, 3, and 4 days in a Petri dish sealed with microporous tape, then co-cultured for two days in a Petri dish sealed with Parafilm, which led to median transformation efficiencies of 0% (mean: 0.6%), 74.1% (mean: 62.6%), 74.1% (mean: 70.3%), 47.4% (mean: 45.8%), and 9.1% (mean: 12.2%), respectively (Fig. 2b, see also Supplementary Table S2). These results indicate that pre-culture periods of one and two days are optimal.

The use of Parafilm-sealed Petri dishes shortened the period required for the AgarTrap co-culture step. For the following investigations, we used fixed conditions of a two-day pre-culture, a two-day co-culture with *Agrobacterium* strain GV2260 in the light in Petri dishes sealed with Parafilm, and no surfactant in the transformation buffer. These conditions were varied as described below, to investigate their impact on transformation efficiency.

### Surfactants in transformation buffer

In previous studies of *Agrobacterium*-mediated transformation, it was reported that the use of surfactants in the co-cultivation medium during co-culture increased the transformation efficiency^27,28^. We therefore examined whether surfactants in the transformation buffer influenced the efficiency of G-AgarTrap.

To determine a suitable surfactant for *M. polymorpha* transformation, a survival test was performed using three surfactants, Silwet L-77, Triton X-100, and Tween 20. We added various concentrations of these surfactants to the transformation buffer and treated the pre-cultured gemmalings with this buffer. After two days of co-culture, the survival rates of the gemmalings were estimated. Four concentrations of Silwet L-77 (0.01%, 0.02%, 0.05%, and 0.1%) were analyzed, resulting in mean survival rates of 100%, 100%, 98.8%, and 11.7%, respectively (Fig. 3a). When 0.01%, 0.02%, 0.05%, or 0.1% Triton X-100 was used, the mean survival rates of the gemmalings were 100%, 100%, 99.2%, and 63.5%, respectively (Fig. 3b). Because gemmalings could not survive in the higher concentrations of Silwet L-77 and Triton X-100, these surfactants may be toxic to *M. polymorpha.* By contrast, when Tween 20 concentrations of 0.01%, 0.02%, 0.05%, and 0.1% were tested, the mean survival rate was 100% for all concentrations (Fig. 3c). Tween 20 seemed to have no effect on gemmaling growth, and was therefore selected for use as a surfactant.

**Fig. 3.**
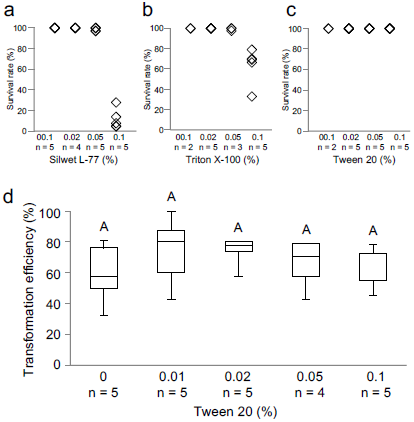
Effect of adding surfactant to the transformation buffer on gemmaling survival rates and transformation efficiency. (a, b, c) The survival rates were estimated for gemmalings treated with various concentrations of Silwet L-77 (a), Triton X-100 (b), and Tween 20 (c). (d) The effect of adding Tween 20 to the transformation buffer on transformation efficiency. All examinations were performed following a two-day co-culture with *Agrobacterium* strain GV2260 under light, in Petri dishes sealed with Parafilm. The same letters indicate no significant difference (Tukey-Kramer’s test; P<0.05).

We assessed whether the use of Tween 20 in the transformation buffer increased the efficiency of G-AgarTrap. Tween 20 concentrations of 0%, 0.01%, 0.02%, 0.05%, and 0.1% resulted in median transformation efficiencies of 57.6% (mean: 59.3%), 80.0% (mean: 74.1%), 77.4% (mean: 73.8%), 70.6% (mean: 65.8%), and 54.8% (mean: 61.1%), respectively (Fig. 3d, see also Supplementary Table S3). These results showed that the use of 0.01–0.02% Tween 20 in the transformation buffer slightly increased the efficiency of G-AgarTrap transformation; however, the differences were not statistically significant. Nevertheless, when the gemmalings were co-cultured in transformation buffer, the solutions lacking surfactant were often repelled by the plants, requiring careful manipulation to ensure proper coverage. When surfactants such as Tween 20 were added to the transformation buffer, this hydrophobicity was counteracted; therefore, the addition of surfactants improves the ease of performing G-AgarTrap transformations.

### *Agrobacterium* strain

*Agrobacterium* strains influence the efficiency of *Agrobacterium*-mediated transformations in other plant species, with the most effective strain being dependent on the plant species or transformation method used^29–33^. For the transformation of *M. polymorpha* above, and in the previous G-AgarTrap study, the GV2260 strain was used^24^. To assess the best strain for G-AgarTrap transformation, we compared the efficiencies of the technique using five *Agrobacterium* strains, GV2260, EHA101, EHA105, LBA4404, and MP90^34–38^. The median transformation efficiencies using these strains were 61.0% (mean: 57.6%), 96.7% (mean: 93.8%), 47.6% (mean: 47.2%), 28.3% (mean: 26.2%), and 9.2% (mean: 18.1%), respectively (Fig. 4a, see also Supplementary Table S4). The use of *Agrobacterium* strain EHA101 resulted in over a 90% efficiency in eight out of 10 transformations, and 100% efficiency on four occasions (Fig. 4a, see also Supplementary Table S4). EHA101 was therefore the superior strain for AgarTrap, contributing consistently high levels of transformation efficiency (Fig. 4a, see also Supplementary Table S4), which also resulted in the presence of many transformed cells within each gemmaling (Fig. 4b). Conversely, MP90 was not suitable for AgarTrap, as its use resulted in a 0% efficiency for two of 10 transformations, and only ever resulted in one or a few transformed cells within a single gemmaling (Fig. 4a, c, see also Supplementary Table S4).

**Fig. 4.**
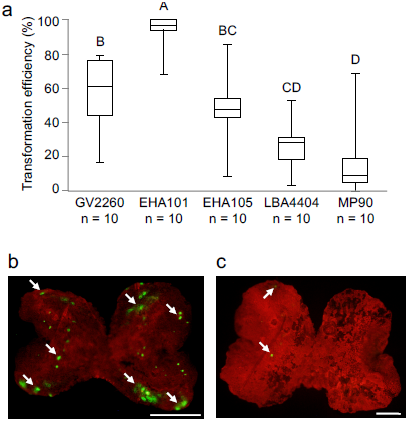
Effect of *Agrobacterium* strain on transformation efficiency. (a) The transformation efficiency of G-AgarTrap using five *Agrobacterium* strains, GV2260, EHA101, EHA105, LBA4404, and MP90. All examinations were performed following a two-day co-culture under light, in Petri dishes sealed with Parafilm. Different letters indicate a significant difference (Tukey’s test; P<0.05). (b, c) Fluorescence images of transient marker expression in a gemmaling transformed using EHA101 (b) and MP90 (c), cultured for three days after treatment with the selection buffer. Red and yellow-green indicate chlorophyll and Citrine fluorescence, respectively. Scale bar, 500 µm. Arrows indicate representative transformed cells.

We assessed the combined use of the most efficient *Agrobacterium* strain, EHA101, and 0.01–0.02% Tween 20 as a surfactant. When gemmalings were transformed with EHA101 in the presence of 0.01% Tween 20, the median transformation efficiency was 95.5% (mean: 93.1%), which was similar to the efficiency of EHA101-mediated transformations without a surfactant (Supplementary Fig. S2, see also Supplementary Table S5). The median transformation efficiency of EHA101 using 0.02% Tween 20 as a surfactant decreased to 17.6% (mean: 40.1%) (Supplementary Fig. S2, see also Supplementary Table S5). Thus, when using EHA101, 0.01% Tween 20 yields better results than 0.02% Tween 20.

### Light/dark condition during co-culture

In previous studies of *Agrobacterium*-mediated transformation, light and dark conditions were reported to influence the transformation efficiency^39–41^. All previous studies of AgarTrap were performed under continuous white light conditions (75 µmol photons m^−2^ s^−1^) ^23–25^. When *M. polymorpha* was co-cultured with *Agrobacterium* strain GV2260, the median transformation efficiencies under light and dark conditions were 61.5% (mean: 61.3%) and 97.1% (mean: 95.3%), respectively (Fig. 5a, see also Supplementary Table S6). Additionally, the combined use of the most efficient *Agrobacterium* strain, EHA101, and dark conditions during the co-culture period resulted in a median transformation efficiency of 100% (mean: 97.0%), which was the highest efficiency observed in this study. Of the seven transformations performed in darkness using EHA101, a transformation efficiency of 100% was achieved five times (Fig. 5a, b, see also Supplementary Table S6). Numerous cells in each gemmaling were transformed under the dark condition when using either GV2260 or EHA101 (Fig. 5c, d). Thus, for the G-AgarTrap transformation of *M. polymorpha*, the transformation efficiency when gemmalings were co-cultured with *Agrobacterium* under dark conditions was higher than that under light conditions.

**Fig. 5.**
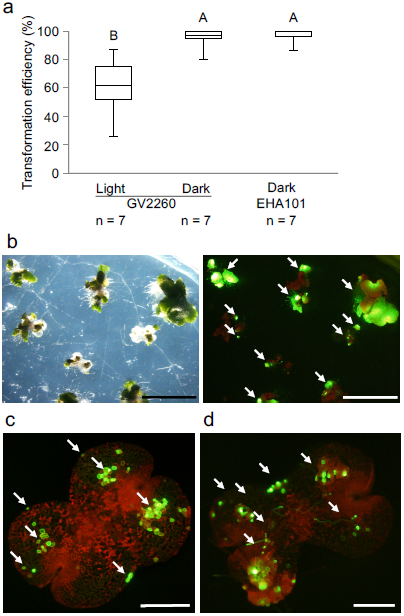
Effect of dark treatment on transformation efficiency. (a) Transformation efficiency following co-culture under light and dark conditions using *Agrobacterium* strain GV2260, or following dark culture using strain EHA101. The effects of the light and dark conditions were examined following a two-day co-culture on Petri dishes sealed with Parafilm. Different letters indicate statistically significant differences (Tukey’s test; P<0.05). (b) Transmitted light image (left) and fluorescence image (right) of stable marker expression in transformants generated using EHA101 in the dark, which were cultured under light for two weeks after treatment with the selection buffer. Scale bar, 0.5 cm. Arrows indicate representative transformants. (c) Fluorescence image of transient marker expression in a gemmaling transformed in darkness using GV2260, and cultured under light for three days after treatment with the selection buffer. (d) Fluorescence image of transient marker expression in a gemmaling transformed in darkness using EHA101, and cultured under light condition for five days after treatment with the selection buffer. For (c) and (d), the scale bar represents 500 µm, red and yellow-green indicate chlorophyll and Citrine fluorescence, respectively, and arrows indicate representative transformed cells.

## Discussion

To improve the efficiency of the G-AgarTrap transformation of *M. polymorpha*, we focused on optimizing the co-culture step for the following four factors: (1) humidity, (2) surfactant in the transformation buffer, (3) *Agrobacterium* strain, and (4) light/dark condition. Among these factors, humidity, *Agrobacterium* strain, and light/dark condition contributed to increases in transformation efficiency.

Because AgarTrap is performed on a solid medium, we predicted that humidity might influence the transformation efficiency. We found that high humidities during co-culture promoted transformation efficiency, and that sealing the Petri dishes with Parafilm instead of microporous tape could overcome the problem of low culture room humidity. The high humidity also enhanced *Agrobacterium* growth, suggesting that this bacterium is sensitive to drying out. Sealing the Petri dishes with Parafilm might better maintain a high internal humidity than sealing the Petri dishes with microporous tape. The enhancement of *Agrobacterium* observed in Petri dishes sealed with Parafilm might promote transformation efficiency; however, the overgrown bacteria were difficult to completely eliminate in the subsequent selection step of G-AgarTrap. When Parafilm was used to seal the Petri dishes during two days of co-culture, efficient pre-culture periods were one and two days. This result was consistent with our previous study using microporous tape-sealed Petri dishes, in which the humidity was approximately 40%^24^. This suggests that the gemmaling cell states arising after 1–2 days of pre-culture might be the most suitable for transformation.

In the *Agrobacterium*-mediated transformation of *Arabidopsis thaliana*, the use of a surfactant, Silwet L-77, increases the transformation efficiency by reducing the surface tension of the aqueous solution^27,42^. In the present study, we did not find any significant improvement in transformation efficiency when using a range of surfactants; however, the addition of surfactants simplified the procedure by reducing the hydrophobicity of the gemmalings, which otherwise repelled the transformation solution. When 0.05% and 0.1% Tween 20 were used, the transformation efficiency using *Agrobacterium* GV2260 was decreased relative to the efficiency when using 0.01% and 0.02% Tween 20 solutions, even though ∼1% Tween 20 did not cause damage to *M. polymorpha* gemmalings. The solutions did not appear to affect the survival rate of *M. polymorpha*; therefore, the higher concentrations (0.05% and 0.1%) of Tween 20 might affect the bacterium itself. The inclusion of Tween 20 when using the more effective EHA101 strain also requires caution, because the transformation efficiency was greatly decreased with a 0.02% concentration of Tween 20 in the transformation solution. EHA101 might therefore be more sensitive to Tween 20 than GV2260.

The transformation efficiency of G-AgarTrap varied significantly with the use of different *Agrobacterium* strains; the strains yielding the highest and lowest efficiencies were EHA101 and MP90, respectively. In a previous study using tomato (*Solanum lycopersicum*), it was suggested that differences in transformation efficiency using different *Agrobacterium* strains was caused by variations in the plant tissue mortality^32^, which might also be the case in the present study. Additionally, for many methods using *Agrobacterium*-mediated plant transformation, the co-culture medium was optimized for transformation, but was also used for the culture of both plant material and *Agrobacterium*. By contrast, in AgarTrap, the co-culture was performed on a solid medium (½ B5 supplemented with 1% sucrose in agar) optimized for the growth of *M. polymorpha*, but not optimized for *Agrobacterium*. Thus, the solid medium might negatively affect *Agrobacterium*, leading to differences in transformation efficiency as a result of differences in the adaptability of the *Agrobacterium* strains to the medium.

It was previously reported that the EHA101 and EHA105 strains are genetically almost identical, as EHA105 was developed by the removal of a kanamycin resistance gene from EHA101^37^; however, in G-AgarTrap, we found a remarkable difference in transformation efficiency when using EHA101 in comparison with EHA105. This might suggest that they are less genetically similar than previously thought. This possibility remains to be investigated.

Previous reports using intact tobacco (*Nicotiana tabacum*) seedlings, *A. thaliana* root segments, and tepary bean (*Phaseolus acutifolius*) calli suggested that light enhanced transformation efficiency^39,41^, but another report using carnation (*Dianthus caryophyllus*) stem explants reported that dark conditions resulted in a higher proportion of transformants^40^. No significant differences in transformation efficiency were observed between light and dark conditions in garlic (*Allium sativum*)^43^. These differences suggest that the effects of light on transformation efficiency might depend on the plant species or tissue used. In the present study, we found that performing the co-culture in darkness significantly enhanced the transformation efficiency. The dark-mediated improvement in transformation efficiency for carnation stem explants was previously suggested to be caused by an increased susceptibility to infection in the etiolated tissue, and/or by an enhanced *Agrobacterium* activation^40^. A subsequent report confirmed that *Agrobacterium* activation is greater in darkness^44^. Plants are more susceptible to infection by pathogens at night, because the reactive oxygen species produced by photosynthesis enhance their resistance to attack^45^. Taken together, we hypothesize that the dark-mediated activation of *Agrobacterium* and the increased susceptibility to infection in the gemmaling cells in darkness result in the observed improvement in transformation efficiency when performing the AgarTrap co-culture in the dark compared with the light condition.

In this study, we successfully developed a highly efficient G-AgarTrap procedure by making several modifications (high humidity, darkness, *Agrobacterium* strain EHA101) to the co-culture step. The improved G-AgarTrap technique will benefit future molecular biology studies of *M. polymorpha*. These improved conditions may also be applicable to other AgarTrap methods (S-and T-AgarTrap). Furthermore, understanding the biological mechanisms underpinning the benefits of these improvements may contribute to the enhancement of the many other transformation technologies using *Agrobacterium* applied to various plant species.

## Methods

### Plant materials and growth conditions

*Marchantia polymorpha* (L.) gemmae/gemmalings of BC3-38, the female line of the third backcross generation created in the crossing of Takaragaike-1 (male line) and Takaragaike-2 (female line), were used in this study. BC3-38 was provided by Dr. Takayuki Kohchi (Kyoto University, Kyoto, Japan). The plants were maintained on half-strength Gamborg’s B5 (½ B5) medium^46,47^ containing 1% agar (BOP; SSK Sales Co., Ltd., Shizuoka, Japan), pH 5.5, in a 90-mm disposable sterile Petri dish. *M. polymorpha* tissues were illuminated with 75 µmol photons m^−2^ s^−1^ continuous white light (FL40SW; NEC Corporation, Tokyo, Japan) in a culture room maintained at around 22°C with air conditioning. The gemmae/gemmalings subjected to G-AgarTrap transformation were obtained from one-to two-month-old thalli.

### G-AgarTrap

The basic procedure of G-AgarTrap was previously reported^24^. Gemmae were sown on approximately 10 mL ½ B5 solid medium (1% agar) supplemented with 1% sucrose, pH 5.5, in a 60-mm disposable sterile Petri dish, and pre-cultured into gemmalings. For the co-culture, 1–3 mL transformation buffer (10 mM MgCl_2_; 10 mM MES-NaOH, pH 5.7; 150 µM acetosyringone; *Agrobacterium* OD_600_ = 0.5) was poured over the gemmalings, with the excess being removed after 1 min using an aspirator or micropipette. Four factors were considered, including sealing of the Petri dish with Parafilm, the *Agrobacterium* strain used, the addition of a surfactant (0.01–0.1% Tween 20) in the transformation buffer, and dark treatment during the co-culture period. After co-cultivation, the *Agrobacterium* was twice washed from the gemmalings and solid medium with 1–4 mL sterile water, and then 1 mL selection buffer containing antibiotics (100 µg hygromycin B and 1 mg Claforan) was poured over the gemmalings and the solid medium. After culturing for a few weeks, the transformed cells had grown and the non-transgenic cells had died^24^.

### *Agrobacterium* preparation for G-AgarTrap

*Agrobacterium tumefaciens* harboring the *pMpGWB103-Citrine* vector, which encodes bacterial aminoglycoside resistance (aadA), was stored in 30% glycerol at –80°C. On the same day that the gemmae were sown on the ½ B5 medium (the first step in the G-AgarTrap procedure), *Agrobacterium* was streaked onto Luria-Bertani (LB) solid medium (1% agar) supplemented with 100 mg L^−1^ spectinomycin and incubated at 28°C for 2–3 days (Supplementary Fig. S1a). The *Agrobacterium* was then suspended in transformation buffer at OD_600_ = 0.5 (Supplementary Fig. S1b). Surfactant (0.01–0.1% Silwet L-77, Triton X-100, or Tween 20) was included in the transformation buffer. A 1-mL aliquot of transformation buffer was poured onto each Petri dish during the co-culture step.

### Microscopy observation

*M. polymorpha* gemmalings were observed using a MZ16F stereo fluorescence microscope (Leica Microsystems, Wetzlar, Germany). Chlorophyll fluorescence and Citrine fluorescence (in transgenic cells) were determined using a fluorescence module (excitation filter: 480/40 nm; barrier filter: LP 510 nm). Images were taken using a DP73 digital camera (Olympus, Tokyo, Japan).

### Transformation efficiency

The transformation efficiency was evaluated using the binary vector *pMpGWB103-Citrine*, which was transformed into *Agrobacterium* as described previously^23–25^. The T-DNA of *pMpGWB103-Citrine* possessed two marker genes encoding hygromycin B phosphotransferase and Citrine fluorescent protein^23–25^. To identify stable transformants, *M. polymorpha* gemmalings were selected for their ability to grow on the antibiotic hygromycin B (10 µg mL^−1^), and their yellow fluorescence was observed using fluorescence microscopy more than two weeks after the selection buffer was poured (transient expression of Citrine has not been observed after this time)^23–25^. A gemmaling containing one or more transformed cells was considered transformed^24^. The transformation efficiency (%) was calculated as the number of transformed gemmalings divided by the total number of gemmalings, multiplied by 100. Approximately 10–50 gemmalings per Petri dish were served for transformation. The median transformation efficiency was considered to be representative, and the mean was also reported to facilitate comparisons with previous studies. Statistics were analyzed by t-test, Tukey’s test, or Tukey-Kramer’s test.

## Acknowledgments

This work was supported by Grant-in-Aid for JSPS Fellows Grant Number 15J09907 (S.T.-T.), the Plant Transgenic Design Initiative of the University of Tsukuba (S.N., H.E., and Y.K.), and the JST-ERATO Numata Organelle Reaction Cluster Grant Number JPMJER1602 (Y.K.).

## Author contributions

S.T., S.N., H.E. and Y.K. designed the research. S.T. and Y.K. wrote the paper. S.T. performed the examinations.

## Additional information

Competing financial interests: The authors declare no competing financial interests.

